# Humidity detection after blood feeding in ticks is required for oviposition site selection and egg survival

**DOI:** 10.64898/2026.11.24.765659

**Authors:** Joshua B. Benoit, Kosisochukwu Onyeagba, Joshua Tompkin, Melissa Kelley, Ronja Frigard, Evan S. Smith, Lyn Wang, Haidar Almatar, Melissa R. Uhran, Brittney Crouse, Kaylee LeVan, Emmarie Watts, Stephanie E. Spitzley, Olivia D. Spencer, Kami M. Fleming, Kylee Brenenborg, Taylor Wood, Derek Herges, Jaishna Sivakumar, Hannah Mahoney, Alekhya Kondragunta, Thomas Arya, Samantha Packwood, Joanna Fetzer, Khwahish Singh, Thomas Brennan, Maher Alhawsawi, Ella Roberts, David Maleh, Joshua Sebastian, Sahr Alisher, Sarah Shamray, Sean Cunningham, Cameron Willey, Robert F. Grimaldi, Vishnu Rajkumar, Amaya Hamilton, Pranav Jayakumar, Crystal Stacey, Jasmine Kennedy, Kennan Oyen, Pia U. Olafson

## Abstract

Immediately following a blood meal, the female tick must locate conditions that allow successful egg development. A contributing factor is that conditions must be sufficiently moist to prevent egg dehydration. Previous studies show that humidity detection occurs through the forelegs and that interference with this process leads to tick mortality; however, this effect has not been examined in fed tick stages. Here, we show that female ticks exhibit a distinct attraction to higher humidity, directing them to locations where they arrest and subsequently deposit eggs. Interference with humidity detection caused fed females to deposit eggs at RHs below the optimal range for egg viability. Evaluation of tick egg viability at cluster sizes of 1, 5, 50, and naturally laid clusters (over 500 eggs) in relation to varying relative humidities reveals that depositing clusters of eggs under higher humidity conditions is critical for viability. Egg viability remained low below 85% RH but increased substantially at higher humidity levels. We found that application of the repellent DEET (*N,N*-diethyl-meta-toluamide) disrupted this behavior. Field validation shows a drastic reduction in surviving larvae when females cannot detect humidity (mechanical or chemical inhibition) during egg deposition. These studies confirm that fed female ticks must detect a humid site after detachment from the host to ensure egg viability, and that disrupting this process could be an effective tick control strategy.

## Introduction

Ticks serve as vectors for numerous disease-causing pathogens, yet they spend most of their lives between blood meals, especially species that require multiple hosts. During these off-host periods, ticks risk dehydration because of their small size and high surface-area-to-volume ratio (Benoit and Denlinger, 2010; Benoit and Oyen, 2021). Maintaining water balance is vital for their survival (Needham and Teel, 1991; Rosendale et al., 2017). To avoid dehydration, ticks employ adaptations to conserve water or to absorb external water (Benoit and Denlinger, 2010; Benoit and Oyen, 2021). One key strategy involves an active, solute-driven process of water vapor absorption from the environment (Gaede and Knülle, 1987; Rosendale et al., 2016). While this process helps replenish their water supply and ultimately prevents death from dehydration (Randolph and Storey, 1999; Rosendale et al., 2017), it also depletes their lipid reserves. Since ticks feed exclusively on blood, their nutrient reserves are limited to those acquired from the egg and subsequent blood meals (Rosendale et al., 2019). Though ticks can endure extended periods of starvation, partly due to their low metabolic rates (Fielden and Lighton, 1996; Lighton and Fielden, 1995), depletion of energy reserves is a significant cause of mortality in the wild (Nieto et al., 2010). Therefore, understanding the factors that enable tick survival between blood meals is crucial for assessing the potential for pathogen transmission.

An important period for tick survival occurs during the egg and larval stages, due to a lack of motility and the exceptionally small size of tick eggs and larvae (Benoit et al., 2007; Yoder et al., 2012a). Although the egg stage has relatively high resistance to dry conditions (Benoit et al., 2007; Yoder et al., 2012a, 2004), prolonged exposure to conditions below 75% RH substantially reduces egg viability (D’Alessandro et al., 2014; Fujimoto, 1990). If eggs are viable, larval survival requires humidities above the critical equilibrium humidity (Gaede and Knülle, 1987; Maldonado-Ruiz et al., 2020; Yoder et al., 2012b). To ensure eggs are deposited in a favorable location, recently fed females must locate a humid microhabitat that maximizes egg viability. The main sensory organ of ticks is the Haller’s organ, a sensory structure on the first pair of legs (Carr et al., 2017). Recent studies indicate that Haller’s organ is directly involved in humidity detection in ticks before feeding (Uhran et al., 2024), but it remains unclear how Haller’s organ and humidity detection influence fed tick behavior, particularly during the off-host search for an optimal oviposition site.

In this study, we tested whether fed female ticks actively use humidity cues to select oviposition sites, whether disrupting Haller’s organ impairs this behavior, how humidity and egg clustering influence egg viability, and whether chemical interference with DEET alters these processes. We further evaluated the potential ecological consequences of disrupted oviposition behavior under semi-field conditions. Haller’s organ interference impaired the ability of fed females to deposit their eggs at optimal relative humidities. When deposited at these suboptimal conditions, egg viability was greatly reduced. Field observations in mesocosms support that the inability to locate an optimal egg-deposition site reduces egg viability. Lastly, gene expression analyses of the forelegs showed high levels of ionotropic receptors associated with humidity detection. Overall, we demonstrate the importance of the Haller’s organ in humidity detection of fed female ticks and in ensuring that tick eggs are deposited under optimal conditions for offspring viability.

## Materials and Methods

### Ticks

Fed female *D. variabilis* were acquired from Ectoservices (Cary, NC), and *Amblyomma americanum* and *D. albipictus* were acquired from the USDA Knipling-Bushland U.S. Livestock Insects Research Laboratory. Ticks from Ectoservices were fed on sheep (*Ovis aries*), and those from the USDA were fed on cattle (*Bos taurus*). Ticks were collected immediately after detachment from the host and held for 12 hours at 93% relative humidity (RH) with saturated salt solutions (Winston and Bates, 1960). Ticks were stored at 24-25 °C and 12:12 h light-dark (L:D) cycle during the course of the assays.

### Interference with the foreleg to impair Haller’s organ function

To evaluate how the Haller’s organ contributes to humidity detection, we conducted a series of manipulations such as wax covering, heat ablation, and leg removal, which are methods previously used in unfed ticks (Uhran et al., 2024). For leg removal treatments, either the first or second pair of legs was removed, allowing comparison between ticks with an intact Haller’s organ and those without. In the heat ablation treatment, a heated metal probe was used to cauterize the Haller’s organ or an equivalent area on the second pair of legs. For the wax covering treatment, the forelegs were coated with warmed dental silicone (Defend), which could later be removed to restore humidity-sensing ability. After all manipulations involving the legs or Haller’s organ, ticks were maintained at 95-100% RH for two days to allow for recovery prior to experimentation. Lastly, DEET (*N,N*-diethyl-meta-toluamide) was applied to the forelegs of ticks to evaluate its effects on humidity detection. Briefly, a 5% DEET solution (diluted in a 95:5 water:ethanol solution) was applied to the fed ticks 30 minutes before the assays by wetting the forelegs with either the DEET or water:ethanol solution with a small brush.

### Humidity preference assays

To evaluate ticks’ ability to detect humidity and initiate quiescent behavior under different moisture conditions, we designed a two-choice humidity assay. A 42-cm polycarbonate tube (2.54 cm i.d.) was fitted at both ends with Drosophila vials (25 × 95 mm) containing materials that produced distinct relative humidity levels: calcium sulfate (Drierite, Xenia, OH) for low humidity (0–10% RH), saturated sodium chloride solutions for medium humidity (75% RH), and sponges soaked in deionized water for high humidity (95–100% RH). Individual ticks were placed in the center of the tube, and each end was capped with a vial containing one of the two humidity conditions being compared, creating a 40-cm attraction distance. Each condition (sponges and desiccant) was refreshed between trials to ensure accuracy and stability of the humidity gradient. Tick positions were recorded after 24 hours, reflecting the location where each tick was most likely to settle for longer quiescent periods. Individuals who did not survive the 24-hour period were excluded from analysis. Along with the position where the tick stopped, the final position where each female deposited eggs in the tubes was recorded. To standardize results, positions were converted to a humidity-attraction value based on distance from the dry end; thus, a tick located 40 cm from the dry side received an attraction value of 1, whereas a tick located directly at the dry side received a value of 0.

### Egg viability

For our study, fed females were delivered to the laboratory within 24–48 hours after host drop-off. Upon arrival, they were maintained at 24-25 ± 1°C, 93% relative humidity (RH), and a 12:12 hr L:D cycle (Winston and Bates 1960). Once oviposition began, eggs were collected and placed in different controlled-humidity conditions. Relative humidities were established in sealed glass desiccators using saturated salt solutions with excess solid salt for 33% (MgCl_2_), 75% RH (NaCl), 85% RH (KCl), and 93% RH (KNO₃) (Winston and Bates 1960); and distilled water for 95-100% RH. Each desiccator’s RH was monitored daily with a hygrometer (Thomas Scientific, Philadelphia, PA, U.S.A.) and fluctuated by only 1–2% during the experiment. To avoid bias, egg batches from multiple feedings were used to complete the studies. Eggs were divided into four groups: individuals, groups of 5-10 (denoted as 5), groups of 50-150 (denoted as 50), and full egg clusters (natural). After eight weeks, the number of hatched larvae and unhatched eggs was counted three times independently by separate researchers to ensure accuracy. Unhatched eggs were typically collapsed at the time of assessment and considered non-viable. Each group of eggs was counted three times by independent observers, yielding 2,811 technical replicates for 937 biological replicates across all treatments for egg viability.

The proportion of eggs that hatched was calculated by dividing the number of hatched eggs by the total number of eggs for each treatment. We tested the relative proportions of eggs that hatched using generalized linear models (GLMs) to compare specific egg groups across varying relative humidities. A generalized linear mixed-effects model (glmer - lme package) was used to evaluate the combined effects of relative humidity and cluster size. For all models, a residual analysis plot (residuals vs. expected values) was visually examined to verify that the model assumptions of normality and homoscedasticity were met. Response variables were logit-transformed as needed to ensure model assumptions were met. Post-hoc comparisons were made using emmeans (default: Tukey method), with p < 0.05 considered statistically significant. All analyses were done in R version 4.3.1, and data visualization was completed using tidyr, ggplot2, and reshape2.

### RNA-seq analyses of winter tick forelegs

RNA-seq analyses were performed as previously described (Benoit et al., 2025; Sterkel et al., 2025) to examine the transcriptional profiles of the first and second pairs of legs in winter ticks. Raw sequencing reads were trimmed using Trimmomatic (version 0.38) (Bolger et al., 2014), and read quality was assessed with FastQC (0.11.5; https://www.bioinformatics.babraham.ac.uk/projects/fastqc/). Trimmed reads were mapped to the *D. albipictus* reference genome using HISAT2 (version 2.1.0). Gene count tables were generated with HTSeq (https://htseq.readthedocs.io/en/latest/) under default parameters. Differential gene expression was determined using the generalized negative binomial model implemented in DESeq2 (version 1.18.1) in R (R Core Team, version 3.5.2). P values were corrected for multiple testing using a false discovery rate (FDR) approach, and genes with adjusted P ≤ 0.05 were considered differentially expressed. Analyses compared transcriptional differences between the first and second set of legs. Following identification of differentially expressed transcripts, enriched functional categories were identified using gProfiler (Raudvere et al., 2019) and summarized with REVIGO (Supek et al., 2011). Datasets have been deposited under the NCBI Bioproject PRJNA1557474.

### Semi-field evaluation of larval recovery following foreleg manipulation and DEET exposure

In addition to laboratory-based survival studies, field assessments were conducted at the University of Cincinnati Center for Field Studies (Harrison, OH) to evaluate egg deposition, larval emergence, and subsequent larval survival following oviposition by engorged adult female ticks under natural environmental conditions. Groups of ten engorged adult female ticks were placed in small field cages (BioQuip, 10 × 10 × 10 cm), which were randomly distributed beneath tree cover to minimize direct exposure to sunlight and provide a shaded environment representative of natural tick habitats. Locally collected organic debris, including leaf litter and other ground materials, was placed at the bottom of each cage to provide a substrate resembling the natural forest floor and to help maintain environmental conditions suitable for oviposition and larval development. Replicate cages were established for each of four experimental treatments: removal of the front pair of legs, removal of the second pair of legs, DEET application, and an untreated control. Each cage contained an engorged adult female. Following treatment, engorged females were allowed to oviposit directly within their respective field cages, where deposited eggs remained exposed to ambient environmental conditions throughout embryonic development. Following egg hatch, emerging larvae were retained within the same cages, allowing assessment of larval emergence and survival under field conditions without additional handling or transfer.

## Results

### Haller’s organ interference impacts humidity detection and the site of egg deposition

When given a choice between low- and high-humidity environments, control ticks with intact forelegs showed a strong attraction to higher relative humidity for their initial movement and for egg deposition (**Fig. 1A–D**). Disruption of the first pair of legs significantly altered this behavior, where all methods of disruption yield significant impairment of humidity detection (P < 0.05 in all cases). Complete removal of leg I eliminated humidity preference, with ticks displaying no consistent attraction to high humidity for *D. variabilis*. Covering the distal tip of leg I with wax similarly abolished attraction to humidity, but this effect was reversible following wax removal, restoring normal humidity preference. Similar effects were noted for *D. variabilis*, *Amblyomma americanum*, and *D. albipictus* when wax covered. Together, these results demonstrate that structures located on the forelegs—most likely the Haller’s organ—are essential for humidity detection and drive behavioral preference for humid microhabitats in fed female ticks.

**Figure 1.**
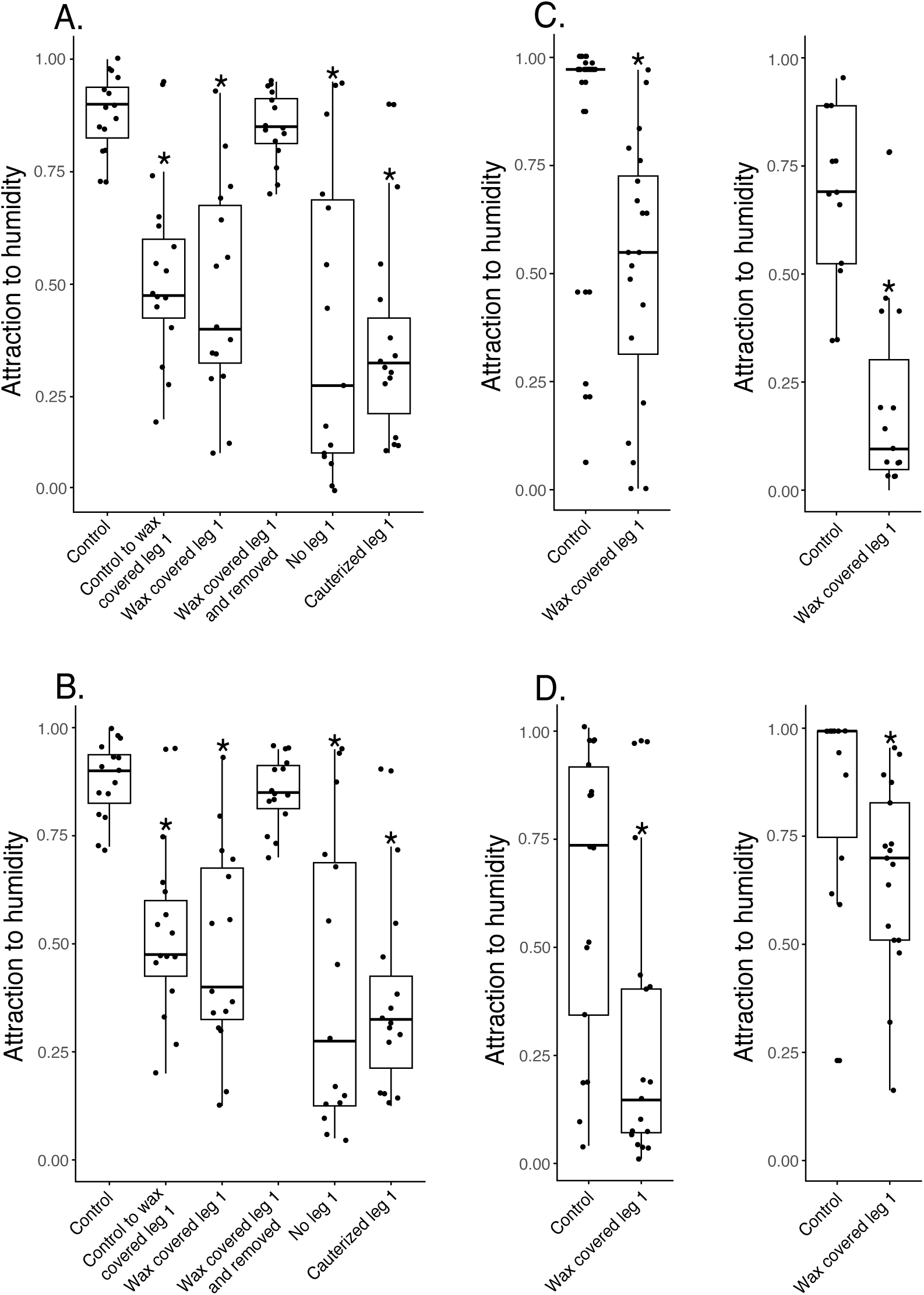
- Haller’s organ on the forelegs is required for humidity attraction in ticks. Attraction to high relative humidity following manipulation of the first pair of legs (leg I) across experimental treatments and species. Attraction to humidity was quantified by the final position of each tick along the humidity gradient after 24 hours, with values normalized to a range of 0 (dry end) to 1 (humid end). Individual data points overlaid on boxplots (median, interquartile range, and full distribution) with N=12-20 per treatment. Control ticks displayed a strong attraction to humid conditions. Covering the distal tip of leg I with wax significantly reduced humidity attraction, while removing the wax restored normal behavior, demonstrating the effect’s reversibility. Complete removal of leg I or cauterization of Haller’s organ similarly impaired humidity preference, indicating that sensory structures located on the forelegs are essential for humidity detection and attraction. Partial manipulations produced intermediate responses, consistent with functional redundancy between forelegs. A and B. American dog tick, *Dermacentor variabilis*, assays 24 hours in the assay and after egg deposition, respectively. C and D Winter tick, *Dermacentor albipictus* (left), and Lone star tick, *Amblyomma americanum* (right), assays 24 hours in the assay and after egg deposition, respectively. Asterisks denote significant differences relative to controls, as determined by generalized linear models. *, denotes significance at P < 0.05.

### RNA-seq of fed female forelegs reveals enrichment for humidity-detecting ionotropic receptors

Principal component analysis (PCA) shows a strong separation between foreleg and second leg for fed female ticks along PC1, which explains 75% of the total variance, while PC2 accounts for 11% variance, indicating that leg identity is the dominant source of transcriptional variation (**Fig. 2A**). These differences yield 1,033 genes with increased expression in the forelegs (**Fig. 2B**). Gene ontology enrichment analyses reveal that differentially expressed genes are strongly associated with multicellular organismal processes, developmental pathways, cell morphogenesis, cell adhesion, and cell projection organization, as well as sensory perception of chemical stimuli and multiple ion transport categories, including sodium and monoatomic ion transport (**Fig. 2C**). As expected, these enriched functional groups suggest structural and sensory differentiation between appendages. Notably, sensory genes associated with humidity detection, the ionotropic receptors Ir25a and Ir93a, are prominently expressed in the forelegs (**Fig. 2D**), indicating environmental-sensing roles. Together, the RNA-seq analysis demonstrates that forelegs exhibit a distinct transcriptional signature, enriched with transcripts associated with humidity detection, in female ticks after a blood meal.

**Figure 2.**
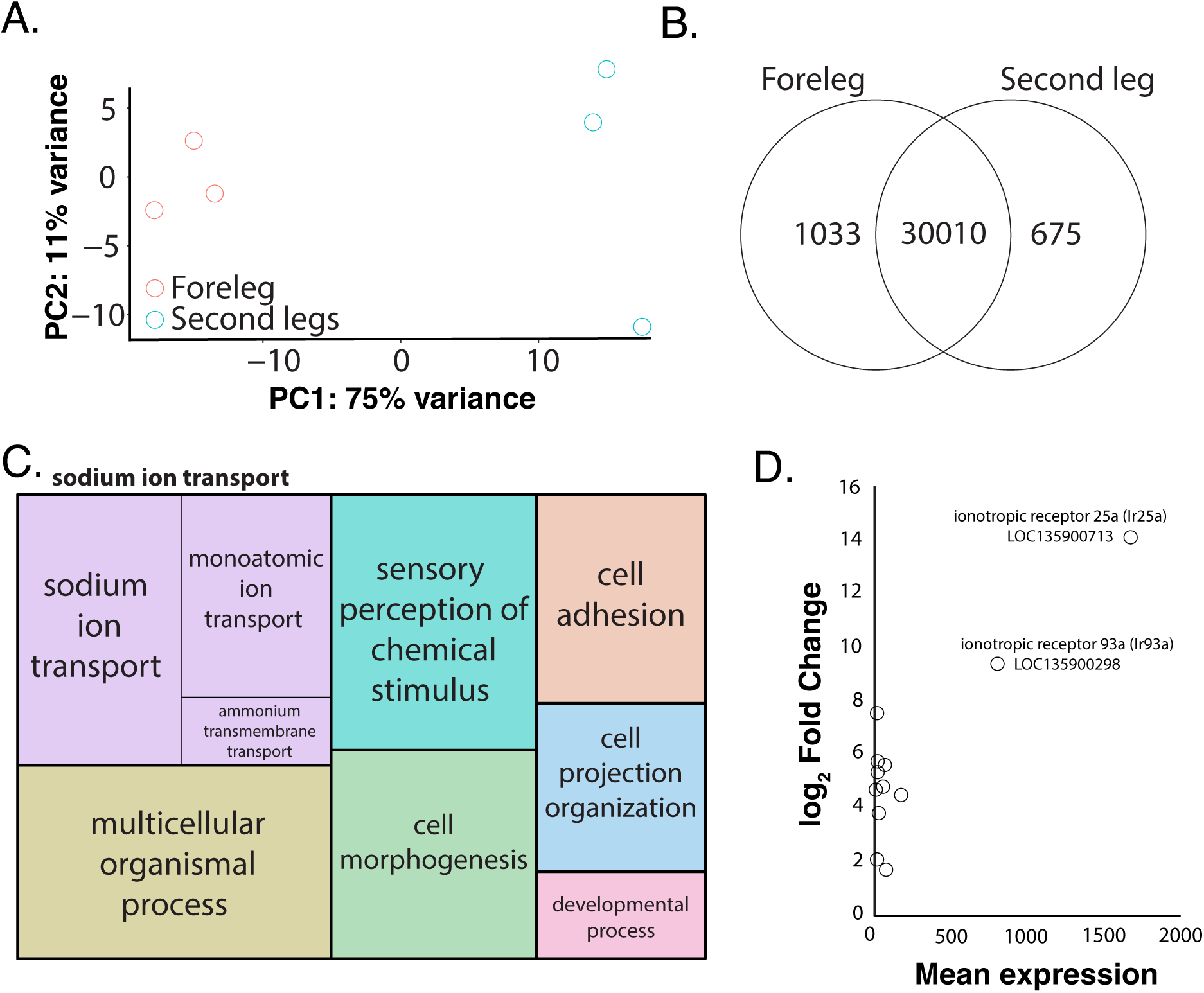
– Transcriptomic differences between forelegs and second legs reveal enrichment of chemosensory pathways in the Haller’s organ location. A. Principal component analysis (PCA) of RNA-seq samples showing clear separation between foreleg and second leg tissues, with PC1 explaining 75% of the variance and PC2 explaining 11% of the variance, indicating strong tissue-specific transcriptional profiles. N = 3. B. Overlap between gene expression in the first and second sets of legs. C. Gene ontology (GO) enrichment analysis of differentially expressed genes, highlighting overrepresented biological processes, including multicellular organismal processes, cell morphogenesis, developmental processes, sensory perception of chemical stimulus, and ion transport pathways. (D) Mean expression versus log₂ fold change plot showing differentially expressed transcripts between forelegs and second legs; key chemosensory-associated genes, including ionotropic receptor 25a (Ir25a) and ionotropic receptor 93a (Ir93a), are highlighted among genes enriched in forelegs.

### Reduced egg viability when females deposit eggs at suboptimal humidities

Egg viability was strongly influenced by relative humidity across all three tick species (*D. variabilis*, *D. albipictus*, and *A. americanum*), with survival increasing consistently as humidity rose. When eggs were tested in standardized groups of 50 (**Fig. 3A**), viability was nearly zero at very low humidity (33% RH) but increased markedly at 75–85% RH and reached the highest levels at 93–100% RH. Although this overall trend was shared across species, sensitivity differed slightly: *D. variabilis* showed a more gradual increase in survival with increasing humidity, whereas *D. albipictus* and *A. americanum* exhibited sharper increases in viability once humidity exceeded moderate thresholds.

**Figure 3.**
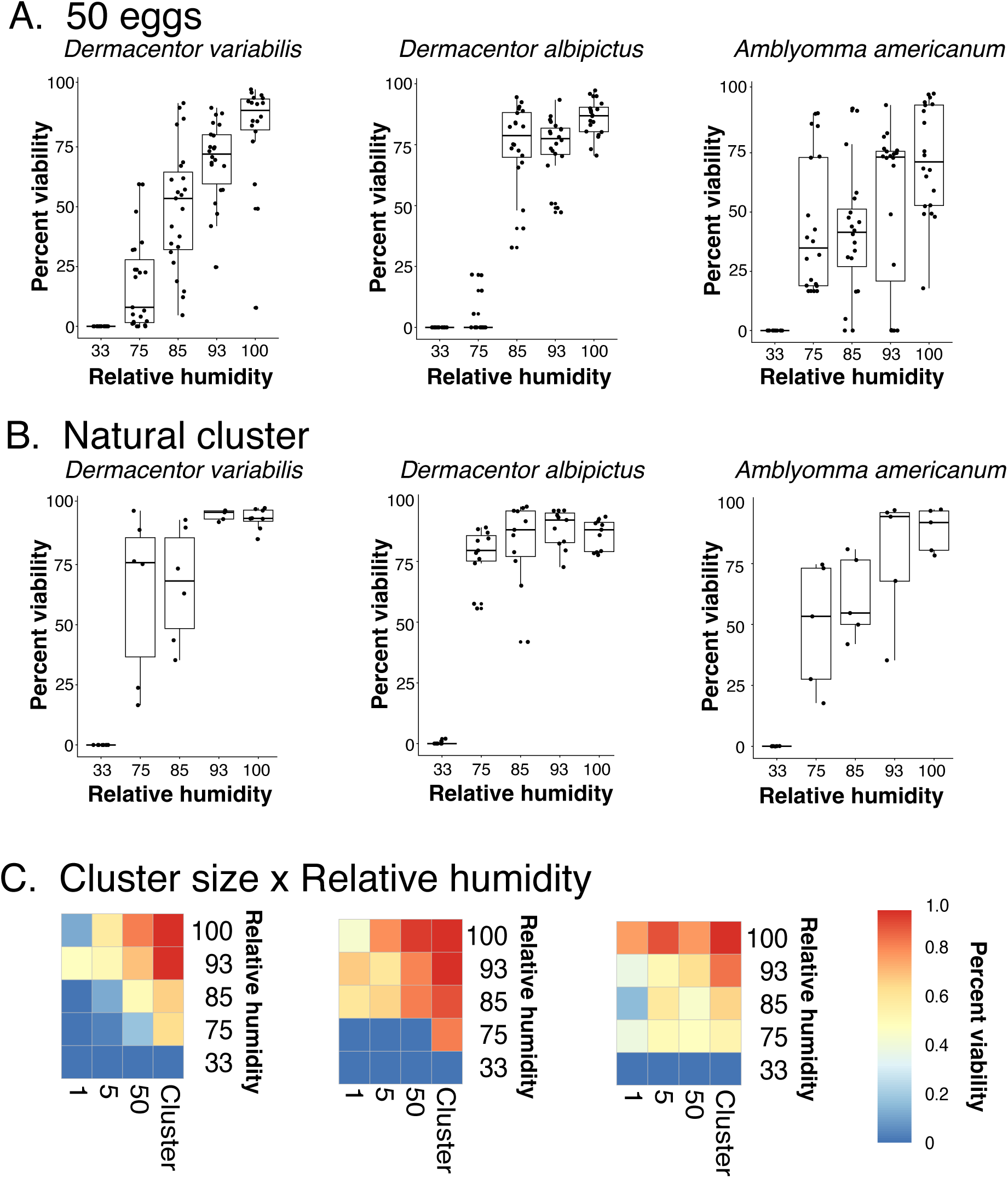
- Effects of relative humidity and egg clustering on tick egg viability across species. A. Percent egg viability for standardized groups of 50 eggs from *Dermacentor variabilis*, *Dermacentor albipictus*, and *Amblyomma americanum* maintained at five relative humidity (RH) conditions (33, 75, 85, 93, and 100% RH). Boxplots show median, interquartile range, and distribution of individual replicates (points). B. Percent viability of eggs maintained in natural oviposition clusters under the same RH treatments for each species, demonstrating enhanced survival compared with standardized egg groups, particularly at intermediate humidities (75-85%). C. Heat maps illustrating the interaction between cluster size (1, 5, and 50 eggs) and RH on egg viability for each species; color scale represents proportional viability (0–1), with warmer colors indicating higher survival. Across species, egg viability increased with humidity and cluster size, indicating that aggregation buffers eggs against desiccation stress. N=8-20 per treatment.

Natural egg clustering substantially improved survival, highlighting the importance of aggregation as a protective strategy against desiccation (**Fig. 3B**). Eggs maintained in natural clusters retained relatively high viability even under intermediate humidity conditions compared with smaller standardized groups. The interaction between cluster size and humidity (**Fig. 3C**) further shows that larger clusters buffered eggs from low-moisture stress, particularly at mid-range humidities (75–85% RH), where survival increased dramatically as cluster size grew from single eggs to large aggregations. Together, these results indicate that both environmental humidity and egg clustering interact to determine reproductive success.

### DEET directly impairs humidity detection

DEET exposure directly limited attraction to humidity in two tick species, *D. albipictus* and *A. americanum* (**Fig. 4**). Both species exhibit reduced attraction to humidity following DEET exposure (P < 0.05 in both cases), with a pronounced decline in *D. albipictus*, where median attraction values drop substantially relative to controls. Although *A. americanum* also shows a decrease in humidity attraction under DEET treatment, the reduction appears more moderate and variable among individuals. The overall trend indicates that DEET interferes with humidity-seeking behavior, suggesting disruption of sensory processes involved in detecting moisture cues.

**Figure 4.**
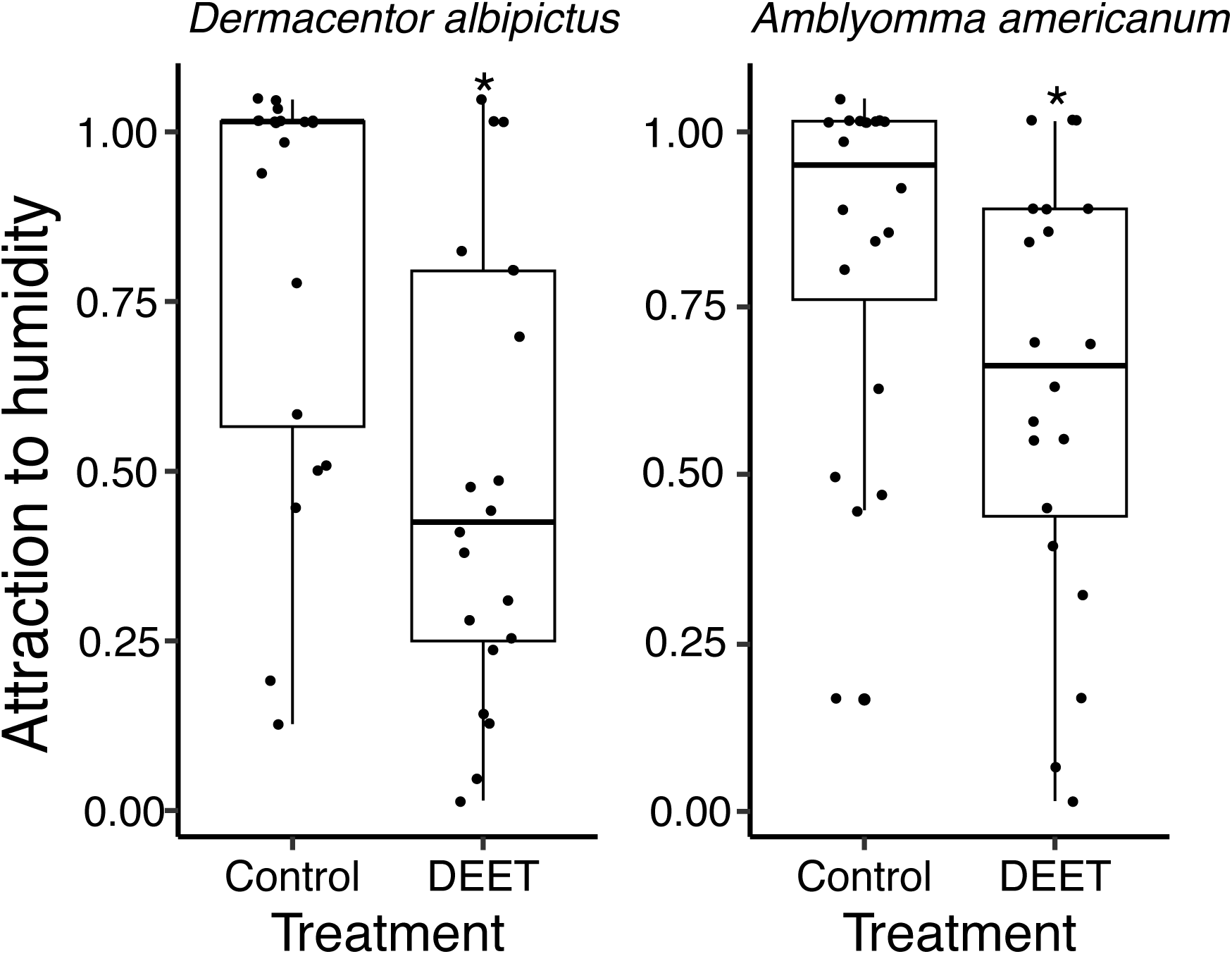
- DEET disrupts humidity attraction in ticks. Attraction to high relative humidity was assessed in *Dermacentor albipictus* and *Amblyomma americanum* following exposure to control conditions or DEET treatment. Boxplots show individual responses (points) with median and interquartile ranges for each treatment group. Control ticks (only exposed to solvent) of both species exhibited strong attraction to humid environments, whereas DEET-treated ticks showed a marked reduction in humidity attraction, indicating that DEET interferes with humidity-guided behavioral responses. *, denotes significance at P < 0.05. N = 12-20 per treatment.

### Larval emergence under suboptimal conditions reduced their survival

Under field conditions, exposure to DEET alters *D. variabilis* tick behavior in a way that reduces the number of larvae collected at the end of the release (**Fig. 5**). Larval recovery is highest in the control condition and substantially lower following DEET treatment and when the first legs were removed (P < 0.05 in both cases). These results indicate that egg and larval survival are reduced under field conditions when humidity detection is compromised.

**Figure 5.**
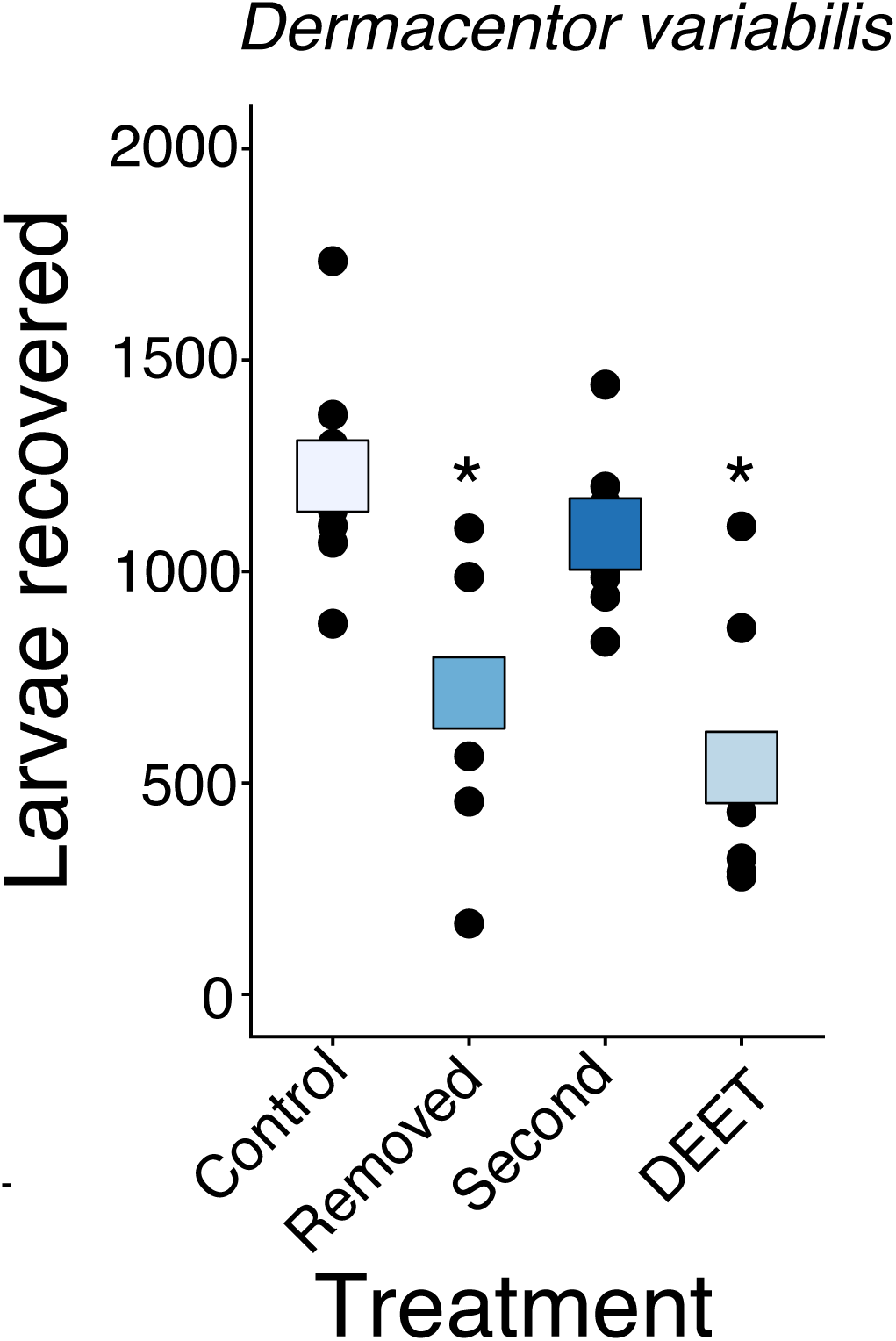
- Effect of foreleg manipulation and DEET exposure on larval recovery in *Dermacentor variabilis*. The number of *D. variabilis* larvae recovered following different treatments is shown. Larval recovery was compared among control ticks, ticks with foreleg (leg 1) or second set (leg 2) removal, and DEET-treated ticks, revealing treatment-dependent differences in larval output. *, denotes significance at P < 0.05. N = 5-6 per treatment.

## Discussion

Our results demonstrate that humidity detection mediated by the Haller’s organ is a primary sensory mechanism guiding oviposition site selection based on humidity in fed female ticks. After detachment from the host, gravid females must locate microhabitats that provide sufficient humidity to support egg development and larval survival. Behavioral assays revealed that control fed ticks with intact forelegs showed a strong attraction to high relative humidity environments, both during initial movement and at the sites where eggs were ultimately deposited. In contrast, interference with the first pair of legs disrupted this behavior. Complete removal of the forelegs abolished humidity preference, whereas unilateral impairment or unilateral cauterization of the Haller’s organ allowed partial recovery of humidity selection, indicating functional redundancy between forelegs. Covering the distal tip of the foreleg with wax similarly eliminated humidity attraction, and removal of the wax restored normal behavior across multiple species. Together, these findings demonstrate that sensory structures located on the tick forelegs—most likely Haller’s organ—are essential for detecting humidity cues that guide gravid females to suitable oviposition environments. These results are similar to those observed in females prior to blood feeding (Uhran et al., 2024), in which humidity detection is required to prevent off-host dehydration.

Transcriptomic analyses further support a specialized sensory role for the forelegs following feeding. RNA-seq comparisons between forelegs and second legs of fed females revealed strong transcriptional separation, with leg identity explaining most variation in gene expression. Over 1,000 genes showed increased expression in the forelegs, with enrichment for developmental and morphogenetic processes, as well as sensory perception and ion transport pathways, which is similar to those previously identified in the forelegs of the black-legged tick, *Ixodes scapularis* (Josek et al., 2018), and in unfed ticks (Uhran et al., 2024). Notably, ionotropic receptors associated with humidity detection in arthropod systems, including Ir25a and Ir93a (Enjin et al., 2016; Knecht et al., 2017; Laursen et al., 2023; Liu et al., 2007), were highly expressed in the forelegs, indicating molecular specialization for humidity sensing after a blood meal. These data provide mechanistic evidence that forelegs function as dedicated sensory appendages for detecting humidity gradients as fed females search for a humidity site to deposit eggs.

Successful oviposition and egg hatching are tightly linked to environmental humidity in ticks (Benoit et al., 2007; Yoder et al., 2012b). Previous egg viability studies in ticks are strongly dependent on high relative humidity, with embryonic development and hatching largely failing under dry conditions. In *Ixodes ovatus*, eggs showed substantial hatching only at very high humidity (≥95% RH), while no hatching occurred at 86% RH or below, demonstrating that even moderate reductions in moisture can prevent successful eclosion (Fujimoto, 1990). Similarly, studies on clustered eggs of multiple tick species found that larval eclosion was maximized at near saturation (>95% RH), whereas low humidity caused dehydration and collapse of the egg chorion, preventing larval emergence and greatly reducing egg viability (D’Alessandro et al., 2014). For our studies, egg viability increased proportionately with relative humidity across all three tick species examined, with survival nearly absent at low humidity but reaching maximal levels under highly humid conditions. Species differed slightly in sensitivity, but all showed strong dependence on elevated humidity for successful embryonic development. Natural egg clustering substantially improved survival as well, demonstrating that aggregation serves as a protective strategy against desiccation, consistent with previous observations (D’Alessandro et al., 2014). Larger egg clusters buffered embryos from moisture stress, particularly at intermediate humidity levels, as we have shown before (Benoit et al., 2007; Yoder et al., 2004). Therefore, both maternal site selection and egg aggregation jointly determine reproductive success. These results highlight that oviposition behavior is not simply a matter of egg placement but a critical decision that directly shapes offspring survival.

Chemical disruption of sensory perception further confirmed the importance of humidity detection for oviposition ecology, demonstrating that DEET could be developed as a novel method to reduce tick levels. DEET is most commonly used as a tick repellent, typically applied to a treated surface or substrate to prevent ticks from contacting the host. Here, we show that direct application of DEET to ticks significantly reduced attraction to humid environments in *D. albipictus* and *A. americanum*, indicating that this compound interferes with sensory pathways involved in moisture detection. While the exact mechanism remains ambiguous, DEET has been shown to interact with sensory receptors (i.e., odorant receptors) in other blood feeding arthropods (DeGennaro, 2015); however, the implications for ionotropic receptors are unclear. The reduction in humidity-seeking behavior suggests that DEET affects environmental sensing beyond host detection, thereby impairing gravid females’ ability to identify suitable oviposition sites. Under field conditions, DEET exposure reduced the number of larvae recovered, demonstrating that disruption of humidity sensing directly reduces recruitment in natural environments.

Collectively, these results establish humidity detection through the Haller’s organ as a central determinant of tick oviposition selection and reproductive fitness. Fed females actively use humidity cues to select oviposition sites that maximize egg viability and larval survival, and disruption of this sensory pathway, whether mechanical or chemical, prevents accurate habitat selection. Transcriptomic evidence reveals molecular specialization of the forelegs for humidity sensing, while behavioral, developmental, and field experiments demonstrate that impaired detection reduces successful recruitment. These findings identify humidity-guided oviposition as a key ecological bottleneck to tick population persistence and suggest that disrupting hygrosensory pathways may provide a novel strategy to reduce tick reproduction and subsequent population growth.

## Acknowledgments

These studies were partially supported through reusable equipment funded by the National Institute of Allergy and Infectious Diseases under Award Numbers R01AI148551, R21AI166633, and R21AI176098 (JBB). The mention of trade names or commercial products is for informational purposes only and does not constitute endorsement or recommendation by the USDA. The USDA is an equal opportunity provider and employer.

## References

Benoit, J.B., Bose, J., Ajayi, O.M., Webster, A., Grieshop, K., Lewis, D., Talbott, H., Polak, M., 2025. Shifted levels of sleep and activity during the night as mechanisms underlying ectoparasite resistance. NPJ Biol Timing Sleep 2, 15.

Benoit, J.B., Denlinger, D.L., 2010. Meeting the challenges of on-host and off-host water balance in blood-feeding arthropods. J. Insect Physiol. 56, 1366–1376.

Benoit, J.B., Yoder, J.A., Lopez-Martinez, G., Elnitsky, M.A., Lee, R.E., Denlinger, D.L., 2007. Habitat requirements of the seabird tick, *Ixodes uriae* (Acari: Ixodidae), from the Antarctic Peninsula in relation to water balance characteristics of eggs, nonfed and engorged stages. J. Comp. Physiol. B 177, 205–215.

Benoit, J., Oyen, K., 2021. Drought and tick dynamics during climate change, in: Climate, Ticks and Disease. CABI, Wallingford, pp. 67–73.

Bolger, A.M., Lohse, M., Usadel, B., 2014. Trimmomatic: a flexible trimmer for Illumina sequence data. Bioinformatics 30, 2114–2120.

Carr, A.L., Mitchell, R.D., III, Dhammi, A., Bissinger, B.W., Sonenshine, D.E., Roe, R.M., 2017. Tick Haller’s organ, a new paradigm for arthropod olfaction: how ticks differ from insects. Int. J. Mol. Sci. 18. 10.3390/ijms18071563

D’Alessandro, W.B., Rodrigues, J., Fernandes, E.K.K., Luz, C., 2014. Impact of humidity on clustered tick eggs. Parasitol. Res. 113, 3899–3902.

DeGennaro, M., 2015. The mysterious multi-modal repellency of DEET. Fly (Austin) 9, 45–51.

Enjin, A., Zaharieva, E.E., Frank, D.D., Mansourian, S., Suh, G.S.B., Gallio, M., Stensmyr, M.C., 2016. Humidity sensing in *Drosophila*. Curr. Biol. 26, 1352–1358.

Fielden, L.J., Lighton, J.R.B., 1996. Effects of water stress and relative humidity on ventilation in the tick *Dermacentor andersoni* (Acari: Ixodidae). Physiol. Zool. 69, 599–617.

Fujimoto, K., 1990. Ecological studies on ixodid ticks : 7. The effects of humidity on oviposition, development and survival of Ixodes ovatus Neumann (Acarina : Ixodidae). Medical Entomology and Zoology 41, 331–339.

Gaede, K., Knülle, W., 1987. Water vapour uptake from the atmosphere and critical equilibrium humidity of a feather mite. Exp. Appl. Acarol. 3, 45–52.

Josek, T., Walden, K.K.O., Allan, B.F., Alleyne, M., Robertson, H.M., 2018. A foreleg transcriptome for *Ixodes scapularis* ticks: Candidates for chemoreceptors and binding proteins that might be expressed in the sensory Haller’s organ. Ticks Tick Borne Dis. 9, 1317–1327.

Knecht, Z.A., Silbering, A.F., Cruz, J., Yang, L., Croset, V., Benton, R., Garrity, P.A., 2017. Ionotropic Receptor-dependent moist and dry cells control hygrosensation in *Drosophila*. Elife 6. 10.7554/eLife.26654

Laursen, W.J., Budelli, G., Tang, R., Chang, E.C., Busby, R., Shankar, S., Gerber, R., Greppi, C., Albuquerque, R., Garrity, P.A., 2023. Humidity sensors that alert mosquitoes to nearby hosts and egg-laying sites. Neuron. 10.1016/j.neuron.2022.12.025

Lighton, J.R.B., Fielden, L.J., 1995. Mass Scaling of Standard Metabolism in Ticks: A Valid Case of Low Metabolic Rates in Sit-and-Wait Strategists. Physiol. Zool. 68, 43–62.

Liu, L., Li, Y., Wang, R., Yin, C., Dong, Q., Hing, H., Kim, C., Welsh, M.J., 2007. *Drosophila* hygrosensation requires the TRP channels water witch and nanchung. Nature 450, 294–298.

Maldonado-Ruiz, L.P., Park, Y., Zurek, L., 2020. Liquid water intake of the lone star tick, *Amblyomma americanum*: Implications for tick survival and management. Sci. Rep. 10, 6000.

Needham, G.R., Teel, P.D., 1991. Off-host physiological ecology of ixodid ticks. Annu. Rev. Entomol. 36, 659–681.

Nieto, N.C., Holmes, E.A., Foley, J.E., 2010. Survival rates of immature *Ixodes pacificus* (Acari: Ixodidae) ticks estimated using field-placed enclosures. J. Vector Ecol. 35, 43–49.

Randolph, S.E., Storey, K., 1999. Impact of microclimate on immature tick-rodent host interactions (Acari: Ixodidae): implications for parasite transmission. J. Med. Entomol. 36, 741–748.

Raudvere, U., Kolberg, L., Kuzmin, I., Arak, T., Adler, P., Peterson, H., Vilo, J., 2019. g:Profiler: a web server for functional enrichment analysis and conversions of gene lists (2019 update). Nucleic Acids Res. 47, W191–W198.

Rosendale, A.J., Dunlevy, M.E., Fieler, A.M., Farrow, D.W., Davies, B., Benoit, J.B., 2017. Dehydration and starvation yield energetic consequences that affect survival of the American dog tick. J. Insect Physiol. 101, 39–46.

Rosendale, A.J., Dunlevy, M.E., McCue, M.D., Benoit, J.B., 2019. Progressive behavioural, physiological and transcriptomic shifts over the course of prolonged starvation in ticks. Mol. Ecol. 28, 49–65.

Rosendale, A.J., Romick-Rosendale, L.E., Watanabe, M., Dunlevy, M.E., Benoit, J.B., 2016. Mechanistic underpinnings of dehydration stress in the American dog tick revealed through RNA-Seq and metabolomics. J. Exp. Biol. 219, 1808–1819.

Sterkel, M., Tompkin, J., Schal, C., Guerra, L.R.M., Pessoa, G.C.D., Oliveira, P.L., Benoit, J.B., 2025. Deployment and transcriptional evaluation of nitisinone, an FDA-approved drug, to control bed bugs. Pest Manag. Sci. 81, 2155–2164.

Supek, F., Bošnjak, M., Škunca, N., Šmuc, T., 2011. REVIGO summarizes and visualizes long lists of gene ontology terms. PLoS One 6, e21800.

Uhran, M.R., Onyeagba, K., Sanderson, S.M., Hoque, S.F., Kelley, M., Smith, E.S., Oyen, K., Benton-Anderson, A., Arya, T., Ledezma, A., Almatar, H., Kennedy, J., Frigard, R., Holmes, C.J., Chen, S.-C., Olafson, P.U., Gaff, H.D., Benoit, J.B., 2024. Impaired humidity sensing reduces tick survival by preventing water homeostasis. Physiology.

Winston, P.W., Bates, D.H., 1960. Saturated solutions for the control of humidity in biological research. Ecology. 10.2307/1931961

Yoder, J.A., Benoit, J.B., Opaluch, A.M., 2004. Water relations in eggs of the lone star tick, *Amblyomma americanum*, with experimental work on the capacity for water vapor absorption. Exp. Appl. Acarol. 33, 235–242.

Yoder, J.A., Hedges, B.Z., Benoit, J.B., 2012a. Water balance of the American dog tick, *Dermacentor variabilis*, throughout its development with comparative observations between field-collected and laboratory-reared ticks. Int. J. Acarology 38, 334–343.

Yoder, J.A., Hedges, B.Z., Benoit, J.B., 2012b. Water balance of the American dog tick, *Dermacentor variabilis*, throughout its development with comparative observations between field-collected and laboratory-reared ticks. Int. J. Acarology 38, 334–343.

